## Supplemental File for "WormSpot: a machine learning-powered viability scoring platform in *C. elegans* for *Candida* pathogenicity studies"

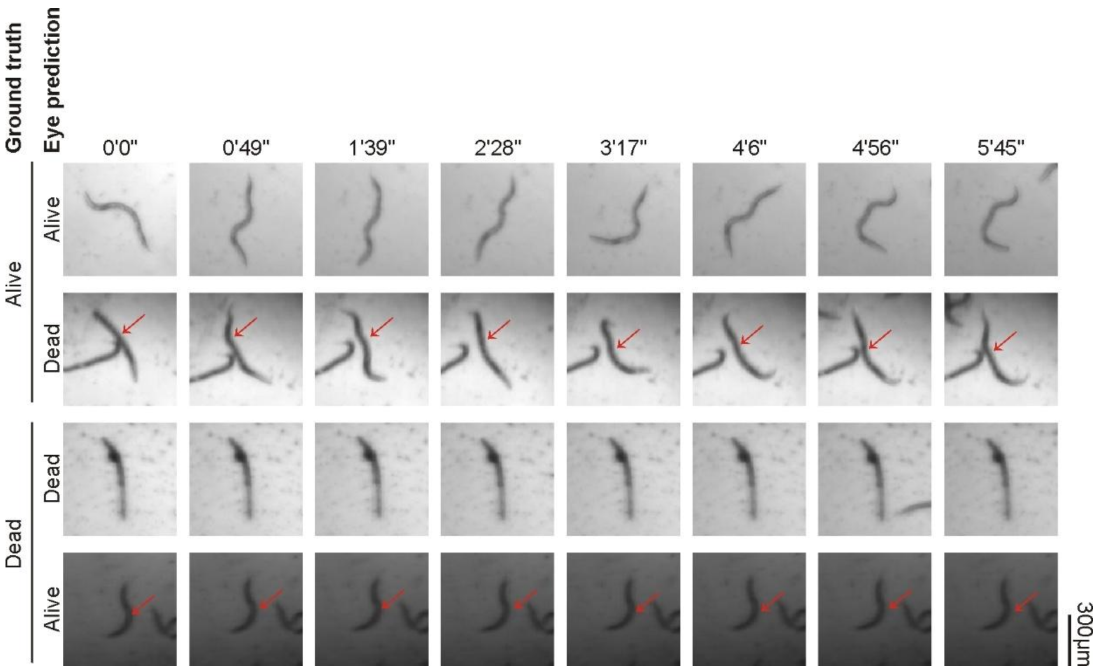


**Figure S1. Video verification of worm viability.**

Example video frames showing correct and incorrect eye predictions of 'Alive' and 'Dead' worms. Red arrows indicate the misclassified worms. Scale bar, 300 μm.

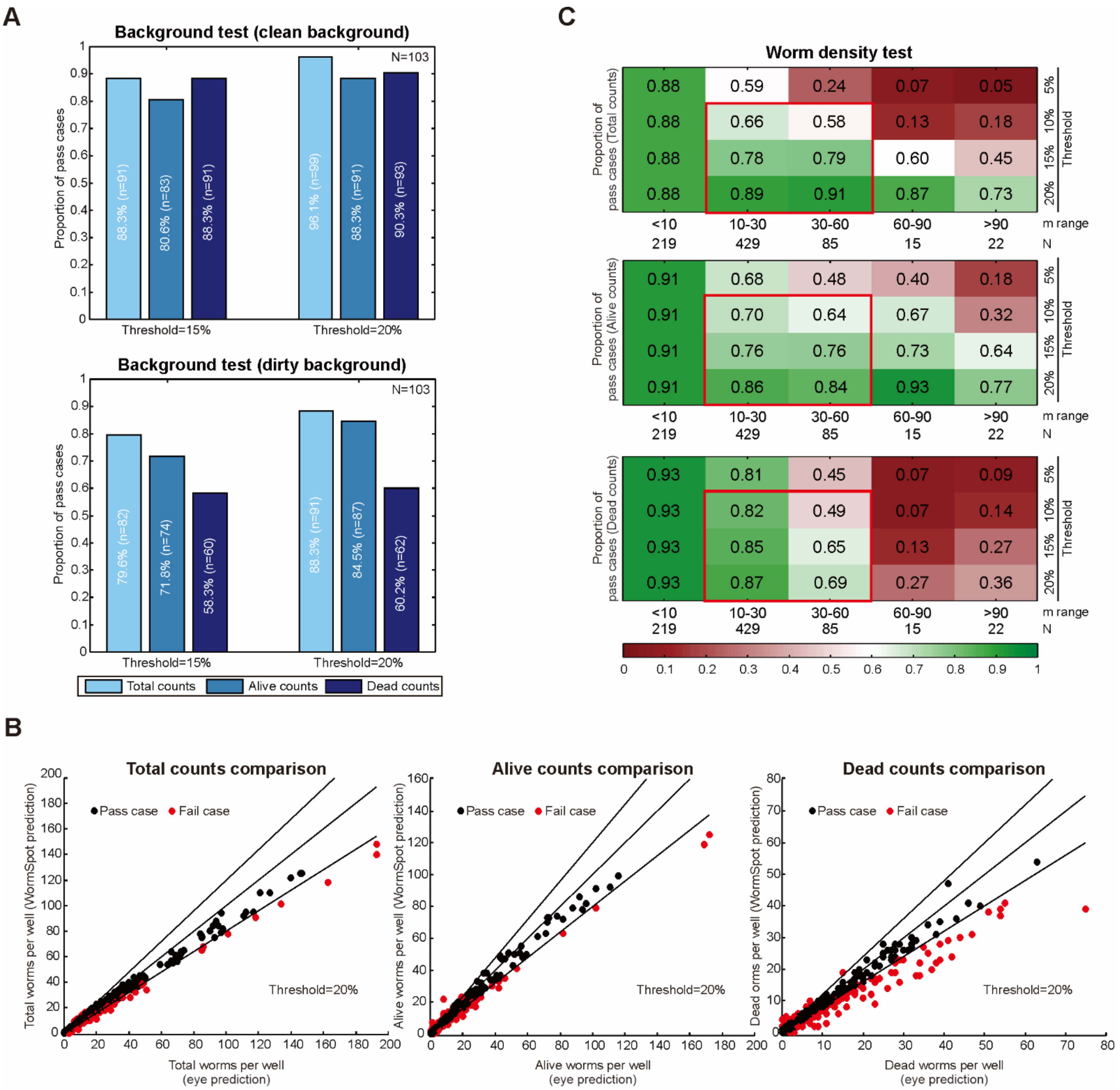


| Proportion of pass cases (Total counts) | <10 | 10-30 | 30-60 | 60-90 | >90 |
|---|---|---|---|---|---|
| 5% | 0.88 | 0.59 | 0.24 | 0.07 | 0.05 |
| 10% | 0.88 | 0.66 | 0.58 | 0.13 | 0.18 |
| 15% | 0.88 | 0.78 | 0.79 | 0.60 | 0.45 |
| 20% | 0.88 | 0.89 | 0.91 | 0.87 | 0.73 |
| N | 219 | 429 | 85 | 15 | 22 |

| Proportion of pass cases (Alive counts) | <10 | 10-30 | 30-60 | 60-90 | >90 |
|---|---|---|---|---|---|
| 5% | 0.91 | 0.68 | 0.48 | 0.40 | 0.18 |
| 10% | 0.91 | 0.70 | 0.64 | 0.67 | 0.32 |
| 15% | 0.91 | 0.76 | 0.76 | 0.73 | 0.64 |
| 20% | 0.91 | 0.86 | 0.84 | 0.93 | 0.77 |
| N | 219 | 429 | 85 | 15 | 22 |

| Proportion of pass cases (Dead counts) | <10 | 10-30 | 30-60 | 60-90 | >90 |
|---|---|---|---|---|---|
| 5% | 0.93 | 0.81 | 0.45 | 0.07 | 0.09 |
| 10% | 0.93 | 0.82 | 0.49 | 0.07 | 0.14 |
| 15% | 0.93 | 0.85 | 0.65 | 0.13 | 0.27 |
| 20% | 0.93 | 0.87 | 0.69 | 0.27 | 0.36 |
| N | 219 | 429 | 85 | 15 | 22 |

**Figure S2. Assessing WormSpot effectiveness for well-based predictions.**

(A) Bar plots separately showing fractions of pass cases for clean and dirty images (N = 103 images each) in the background test at 15% and 20% tolerance thresholds for indicated counts. N, total number of images; n, number of pass cases. (B) Graphs comparing WormSpot predictions (Y-axis) with eye predictions (X-axis) for ‘Total’, ‘Alive’, and ‘Dead’ worm counts in the worm density test. The diagonal line represents exact matches, while the flanking lines indicate 20% threshold boundaries. Dots within the threshold are shown in black; those outside are shown in red. (C) Heatmap showing fractions of pass cases for indicated counts in the worm density test across five specified m ranges under four tolerance thresholds (5%, 10%, 15%, 20%). m, eye-annotated worm count per well; N, image number per m range.

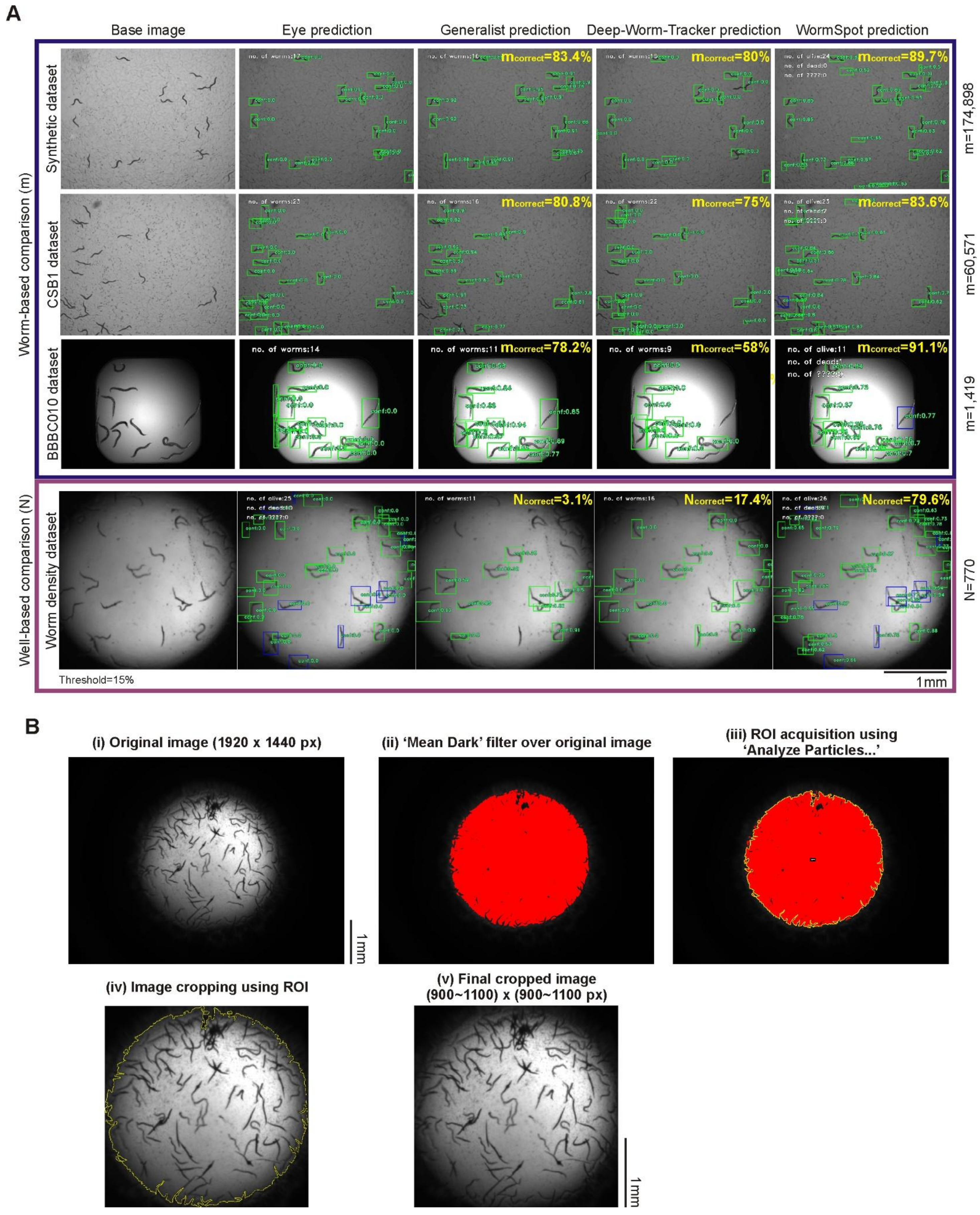
A
Base image
Eye prediction
Generalist prediction
Deep-Worm-Tracker prediction
WormSpot prediction
Worm-based comparison (m)
Synthetic dataset
CSB1 dataset
BBBC010 dataset
m_correct=83.4%
m_correct=80%
m_correct=89.7%
m_correct=80.8%
m_correct=75%
m_correct=83.6%
m_correct=78.2%
m_correct=58%
m_correct=91.1%
m=174,898
m=60,571
m=1,419
Well-based comparison (N)
Worm density dataset
N_correct=3.1%
N_correct=17.4%
N_correct=79.6%
N=770
Threshold=15%
1mm
B
(i) Original image (1920 x 1440 px)
(ii) 'Mean Dark' filter over original image
(iii) ROI acquisition using 'Analyze Particles...'
(iv) Image cropping using ROI
(v) Final cropped image (900~1100) x (900~1100 px)
1mm

**Figure S3. Comparison of worm detection performance between WormSpot, Generalist, and Deep-Worm-Tracker.**
(A) Comparison of worm-based (blue frame) and well-based (purple frame) worm detection performance across WormSpot, Generalist and Deep-Worm-Tracker. Worm-based comparisons used indicated public image datasets, while well-based comparisons used our worm density test dataset at a 15% tolerance threshold. Example images and their corresponding eye annotations and model predictions are shown. Eye-annotated total worm counts (m) for each dataset or well/image counts (N) for the worm density test dataset are listed on the right. We define $m_{correct}$ as the percentage of worms correctly detected and $N_{correct}$ as the percentage of 'pass' cases at the 15% tolerance threshold. (B) Example images showing ImageJ processing steps for generating the final cropped images used in training and testing. Scale bar, 1 mm.
